# The deep-sea ecosystem engineer *Geodia barretti* (Porifera, Demospongiae) maintains basic physiological functions under simulated future ocean pH and temperature conditions

**DOI:** 10.1101/2024.01.11.575017

**Authors:** Erik Wurz, Tone Ulvatn, Tjitske Kooistra, Florian Strauss, Hans Tore Rapp, Jasper M. de Goeij, Ronald Osinga

**Author notes:** author deceased early 2020 (Hans Tore Rapp).

## Abstract

Global ocean warming and acidification will alter the physicochemical conditions in the deep North-Atlantic Ocean. Here, extensive sponge grounds, often dominated by the demosponge species *Geodia barretti*, provide three-dimensional structure, habitat and significantly contribute to benthic-pelagic coupling and nutrient cycling processes in the deep sea. It is unknown if *G. barretti* remains physiologically functional under the future physicochemical properties of an Anthropocene ocean. In this study, individuals of *G. barretti* collected from 300 m water depth in the Barents Sea, were exposed to four treatments resembling future ocean conditions (no treatment, 4 °C increase in seawater temperature, decrease of seawater pH by 0.3, and a combination of the high temperature, low pH). Over the course of 39 weeks, oxygen consumption, dissolved inorganic nutrient fluxes, and bacterioplankton clearance rates were measured as indicators of metabolic activity. We found that all indicators within each sponge individual and per treatment were highly variable over time and no effect of manipulated seawater treatments on these parameters could be demonstrated. Oxygen consumption rates in all groups closely followed a seasonal pattern, potentially caused by (a)biotic cues in the seawater flowing through the experimental aquaria. While similar metabolic rates across all treatments suggest that *G. barretti* physiologically coped with simulated future ocean conditions, observed tissue necrosis in experimental animals might indicate that the response of the complex, high microbial *G. barretti* sponge (i.e., sponge host and microbial symbionts) to future ocean conditions may not be reflected in basic physiological processes.

## Introduction

The North Atlantic Ocean (NAO) harbours vast assemblages of sponges (Porifera) (Cárdenas and Rapp, 2015; Hawkes *et al*., 2019). In these so called deep-sea sponge grounds, sponges can represent up to 98 % of the benthic biomass (Hogg *et al*., 2010) under specific oceanographic settings (Beazley *et al*., 2018; Dijkstra *et al*., 2021; Hanz *et al*., 2021; Roberts *et al*., 2021). These dense assemblages of sponges form complex three-dimensional structures, otherwise scarce in the deep sea, that provide habitat and increase local biodiversity (Beazley *et al*., 2013; Hawkes *et al*., 2019). Often, economically important fish species are associated with sponge grounds (Kenchington, Power and Koen-Alonso, 2013; Kazanidis *et al*., 2018), highlighting their ecological and economic importance. In addition to their role as ecosystem engineers, sponges are key-players in nutrient cycling by filtering large volumes of seawater, removing particulate and dissolved organic matter and regenerate inorganic nutrients (Maldonado *et al.,* 2012(De Goeij et al., 2017). In a confined area (300 km^2^) on the Norwegian shelf the standing stock of the deep-sea sponge species *Geodia barretti* has been estimated to consume 60 t of particulate organic carbon daily (Kutti, Bannister and Fosså, 2013). These magnitudes of organic carbon removal indicate that *G. barretti* dominated sponge grounds recycle organic carbon in the same order of magnitude as other marine invertebrate dominated habitats (Coppari, Zanella and Rossi, 2019). Moreover, in tropical coral-reef ecosystems sponges have been identified as crucial recyclers of dissolved organic matter (DOM) through the so-called sponge loop (De Goeij *et al*., 2013). Recently, also dominant deep-sea sponges, including *G. barretti*, were shown to be able to feed on DOM (Martijn C Bart *et al*., 2020) and a deep-sea sponge loop has been identified (Rix *et al*., 2016; Bart *et al*., 2021). Given that the deep sea represents the largest habitat on the planet (approximated at 360 million km^2^, (Danovaro et al., 2014)) and contains a high abundance of sponges, sponge-driven processes have the potential to play a crucial role in providing global ecosystem services (Armstrong *et al*., 2019) like habitat provision and carbon and nutrient cycling (Ottaviani, 2020; Rossi and Rizzo, 2020). However, to date it is unknown if deep-sea sponges can maintain their ecological function under future changing ocean conditions.

Human-induced climate change is altering the physicochemical seawater properties of the deep NAO (Sweetman *et al*., 2017) and *Geodia*-dominated sponge grounds in the South-Western Barents Sea have been identified as areas impacted by ocean acidification (OA) and ocean warming (OW)(Kazanidis *et al*., 2019). By the year 2100, the water masses in the deep NAO are expected to experience a decrease of seawater pH of up to 0.3 units and a temperature increase of up to 4 °C (Sweetman *et al*., 2017). While tropical shallow water reefs and their sponge fauna have received extensive scientific attention over the past decades (Carballo, J. L., & Bell, J. J. (2017). Climate change, 2017), knowledge about the physiological responses of deep-sea sponges to future ocean conditions is scarce and limited to a few species. *Geodia barretti*, one of the most abundant and well-studied *Geodia* species in sponge grounds in the NAO has been shown to express an increased mortality under heat wave events (temperatures of 4 °C above average) (Guihen, White and Lundälv, 2014), highlighting the sensitivity of cold-water adapted sponge species to changes in environmental conditions (Strand *et al*., 2017). The interaction of OW and OA has been found to have adverse effects on the pumping capacity and tissue integrity in a deep-sea glass sponge from the North East Pacific (Stevenson *et al*., 2020). However, information is lacking about the physiological response of *Geodia* to (combined) near-future ocean conditions and the implications for sponge-driven ecosystem services. Given the large impact climate change will have on all vulnerable deep-sea habitats in the NAO (Johnson *et al.,* 2018), and the significance of sponge-driven ecosystem services in the deep sea, it is crucial to understand how basic physiological functions of highly abundant sponge species like *G. barretti* will respond to near-future ocean conditions.

Oxygen consumption, the clearance of bacterioplankton, and fluxes of inorganic nutrients (e.g., ammonium, nitrate, phosphate) have been used extensively in assessing the metabolic or physiological function of deep-sea sponges, including *G. barretti* (Leys et al., 2018; Bart et al., 2020a, 2021). In *G. barretti*, the pumping of water is directly linked to the retention of bacterioplankton due to the lack of bypasses that would shunt water through the sponge holobiont without being filtered (Leys *et al*., 2018b). Thus, the clearance rate for bacterioplankton can be used as a proxy for the volume of water processed. To cope with the constraints of a warmer, more acidic future ocean, it is of pivotal importance that *G. barretti* can maintain its basic metabolic functions under the predicted future physicochemical seawater conditions. This study is the first to assess the long-term acclimatisation potential of *G. barretti* to near-future sea water pH (decrease by 0.3 units) and temperature conditions (increase of 4 °C). In an aquaria-based, multifactorial experiment, the basic physiological functions of oxygen consumption and bacterioplankton clearance will be assessed over the course of ten months. The assessment of oxygen consumption rates and bacterioplankton clearance rates under manipulated seawater parameters will provide baseline information about the resilience of sponge-driven benthic-pelagic coupling mechanisms in a future ocean. Additionally, the multifactorial design applied in this study will allow for the evaluation of single and combined stressor effects to better understand the effect of (a combination of) future climate effects on our model sponge.

## Materials and methods

### Sponge collection

Individuals of *G. barretti* with a maximum diameter of ∼10 cm were collected by the remotely operated vehicle (ROV) ÆGIR6000 deployed from the Norwegian research vessel GO Sars in the area Tromsøflaket East (71°35’20.4“N 21°22’35.4”E) at a water depth of 330 m. Only undamaged individuals were stored in the collection box and brought to the surface. During sampling at the seafloor, re-suspension of sediment was kept at a minimum. At the surface, sponges stayed submerged in seawater at all times in the ROV’s collection box until they were transferred to the flow-through aquaria facilities onboard GO Sars. Temperature controlled (7 °C, 20 L min^-1^) seawater from 6 m depth was pumped into 40-L tanks on board in which the sponges were kept in the dark throughout the expedition, after which the sponges were transferred to the running seawater aquarium facilities of the University of Bergen, Norway.

### pH and temperature treatments

Sponges were kept in twelve 40-L tanks (Figure 1; 2-3 sponges per tank, 31 sponge individuals in total), supplied with sand-bed filtered seawater obtained from 200 m water depth from the adjacent fjord at a flow rate of 1 L min^-1^. No additional food was added. Three replicate tanks were assigned to one of the following experimental treatment groups: Control (7 sponges), low pH (9 sponges), high temperature (T) (7 sponges) and low pH combined with high T (8 sponges). Seawater manipulation (temperature increase by 4 °C, pH decrease by 0.3 units) was achieved via computer-controlled (Evolution Controller, Aquatronica, Italy) heating elements and the dosing of pressurized CO_2_ (Riebesell, Fabry and Hansson, 2010). pH electrodes (acq310n-ph, Aquatronica, Italy) in the experimental units were triggering low-pressure solenoid valves that opened and closed, when respectively, the upper (pH 8.1) or lower (pH 7.8) setpoint was reached. Sensors measuring seawater pH were cleaned and calibrated weekly following manufacturer guidelines (two-point calibration with pH = 7.00 and pH = 10.00, Certipur^®^, Merk, Germany). Temperature was increased from ambient to treatment levels by 0.6 °C 24 h^-1^, whereas seawater pH was lowered by 0.2 units 24 h^-1^ until reaching treatment levels. An overview of the treatment-specific seawater parameters over the course of the 10-month long experiment can be found in Table 1. Throughout the experiment sponges were visually monitored for signs of tissue necrosis. Tissue necrosis in *G. barretti* was characterized by black spots on the sponge’s cortex that were eventually covered in white, flocculent material (Luter *et al*., 2017). Sponges with signs of necrosis were removed from the experiment to not impact other individuals within the same tank.

**Figure 1.**
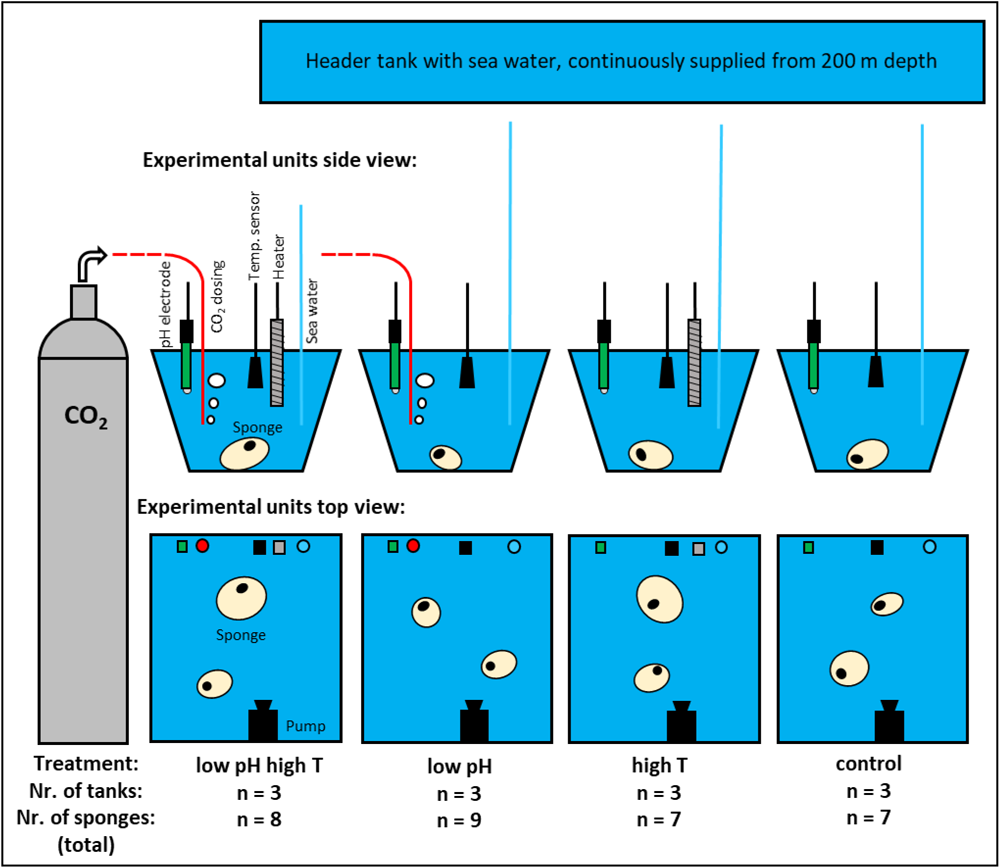
Experimental design used in this study. Individuals of *G. barretti* were exposed to four different sea water treatments. Each treatment had three replicate tanks holding up to three sponges. The tanks in the treatments were individually supplied with seawater from the nearby fjord from a depth of 200 m. Each tank was equipped with a pH electrode and temperature sensor connected to a computer. The treatments high T, low pH and low pH high T were equipped with a heater, CO_2_ dosing, and combination of those, respectively. The heater and CO_2_ dosing were controlled by a computer to maintain treatment parameters. An aquarium pump in each tank provided circulation and mixing to avoid temperature and pH gradients within the tank.

**Table 1.**
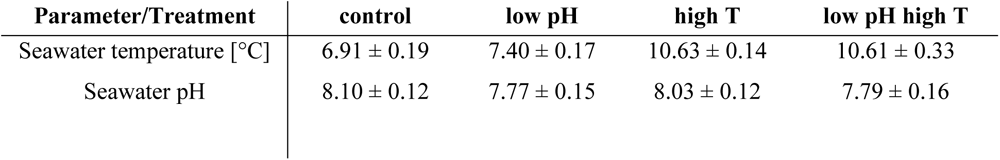
Overview of the sea water parameters in the experimental tanks. Values represent averages over time (39 weeks) and the three tanks per treatment. Values given ± Standard deviation.

### Metabolic rate measurements

Oxygen consumption, bacterioplankton clearance, and inorganic nutrient uptake or release rates of *G. barretti* in the four treatment groups were assessed by chamber-based incubations after 3, 11, 25, 28, 31, and 39 weeks (note: inorganic nutrient concentrations were only assessed in week 11). Therefore, within its respective tank, a sponge individual was carefully manoeuvred onto the bottom plate of a respiration chamber (volume = 6.2 L). Sponges were given a 3 h recovery time before a transparent cylinder and a lid equipped with an optical oxygen probe, and magnetic stirrer for constant water mixing, were fitted onto the bottom plate complementing the air-tight incubation chamber. Individuals were incubated for 4 h.

### Oxygen consumption rates

Concentrations of dissolved oxygen (O_2_) were assessed min^-1^ with an optical sensor (HQ30D53315000, Hach, Colorado, USA) and oxygen removal rates in µmol O_2_ g DM^-1^ h^-1^ were calculated following (Tjensvoll *et al*., 2013) using the following equation: 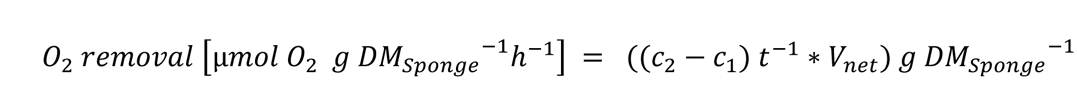, where c_1_ and c_2_ = the concentration of dissolved oxygen at start and end of the incubation in µmol O_2_ L^-1^, t = the time of the incubation in h, V_net_ = the volume of the incubation chamber after subtracting the volume of the respective sponge in mL and g DM = the total dry mass of the incubated animal in g. Sponge mass was assessed at the end of the experiment by taking the respective sponge out of the water and let water drip from the sponge for 10 s before placement on the scale. To determine the dry mass for each individual, per sponge three 1 cm^3^ tissue samples containing cortex and mesohyl were extracted randomly, weighted, dried for 12 h at 60 °C and weighted again. Sponge volumes were assessed at the end of the experiment by measuring the volume of replaced seawater by each individual.

### Bacterioplankton clearance rates

Duplicate water samples (2 mL) were drawn from the chamber at the beginning and the end of the 4-h incubations. Samples for bacterioplankton quantification were fixed with glutaraldehyde (final concentration 0.5 %) for 10 min before flash freezing in liquid nitrogen and −80 °C storage until further analysis. Bacterioplankton abundances were assessed by flow cytometry (Attune NxT – Acoustic focusing cytometer, Invitrogen, Thermo Fisher, USA). Therefore, 500 µL TE buffer were stained with 5 µL of a 1 × working solution of SYBR® green I (Invitrogen, Thermo Fisher, USA) and kept in the dark for 10 min at room temperature. 40 µL of the stained TE buffer were analysed with a flow speed of 25 µL min^-1^. Then, seawater samples (50 µL) were diluted in 450 µL TE buffer and stained with 5 µL of a 1 × working solution of SYBR® green I. After 10 min in the dark at room temperature, 40 µL of stained seawater sample were analysed with a flow speed of 25 µL min^-1^ while keeping the rate of events s^-1^ below 500 to avoid clogging of the machine. Gates were manually set around stained cells that were detected when compared against the stained TE buffer. Clearance rates for bacterioplankton in mL mL_Sponge-1_ h^1^ were calculated as outlined by (Robertson et al., 2017) using the following equation:

### 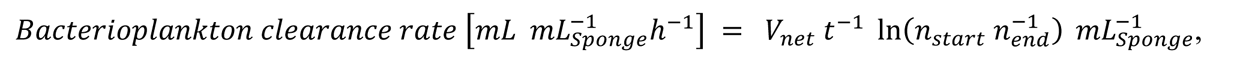

where V_net_ = the volume of the incubation chamber after subtracting the volume of the respective sponge in mL, t = the time of the incubation in h, n_start_ and n_end_ = the concentrations of bacterioplankton at start and end of the incubation in counts mL^-1^ and mL_Sponge_ = the volume of the incubated sponge individual in mL. Seawater-only incubations were performed (n = 4 in weeks 11 and 39, n = 3 in weeks 3, 25, 28 and 31) to account for background respiration and non-sponge related dynamics in bacterioplankton. Sponge incubations were corrected for seawater-only chamber dynamics in oxygen consumption and bacterioplankton concentration. For the correction the average of the seawater-only chamber incubations per timepoint was used.

### Dissolved inorganic nutrient fluxes

Duplicate water samples (10 mL) were taken at the beginning (t_0_) and end (t_2_) of the 4-h incubation to assess the removal and release rates of dissolved inorganic nutrients (ammonium: NH_4+_, nitrate: NO_3-_, nitrite: NO_2-_, and phosphate: PO_43-_) by *G. barretti* (only in week 11). Water samples were filtered over a sterile 0.2 µm polyether sulfone filter (Puradisc, Whatman, UK), collected in high-density polyethylene vial (Pony vial, Perkin Elmer, USA) and stored at −20 °C until further analysis. Concentrations of inorganic nutrients were detected with an automated Wet Chemistry Analyzer (SAN++, Skalar Analytical, Breda, The Netherlands). Nitrate concentrations were derived as: [NO_3-_] = [NO_x−_] – [NO_2−_]. Exchange rates, normalized to sponge dry mass and given in µmol g DM^-1^ h^-1^, were calculated from the difference in concentration over time and the chamber volume, corrected by changes in a control chamber incubation without a sponge.

### Data analysis

Physiological rates of individual sponges were averaged per experimental unit (tank), resulting in *n* = 3 per treatment. The datasets of respiration and clearance rates passed the Shapiro-Wilk test for normality. To investigate the differences between the treatments, we tested for significance at the 5 % level by using a repeated measure, mixed-model analysis of variance with the factors time, temperature and pH. To investigate the relations between individual timepoints, Tukey’s multiple comparisons were performed. When time was not found to have a significant effect, timepoints were averaged and a two-way ANOVA with the factors temperature and pH with Tukey’s multiple comparisons was performed. Treatment effects on nutrient uptake and excretion rates were assessed using dissolved inorganic nutrient specific two-way ANOVAs. Rates of tissue necrosis in *G. barretti* individuals from the four different treatment groups were analysed by a log-rank (Mantel-Cox) test. All tests were performed with GraphPad Prism 8 Version 8.2.1(441), August 20, 2019. Values are given in mean ± standard deviation (SD) if not stated otherwise.

## Results

### Metabolic rates under different pH and T treatments

#### Oxygen consumption rates

Sponges in the control group expressed a trend (*p* = 0.09; PROVIDE STATS) of increased oxygen consumption over time (Figure 2A) ranging from 0.65 ± 0.21 µmol O_2_ gDM^-1^ h^-1^ in Week 3 (time of the year: November; mean ± SD unless stated otherwise) to 1.36 ± 0.11 µmol O_2_ g DM^-1^ h^-1^ in Week 28 (time of the year: July) before returning to levels observed at the start of the experiment in Week 39 (time of the year: September). Oxygen consumption rates were not affected by manipulated seawater conditions (Temperature: *p* = 0.07, pH = 0.92, Table 3) in the respective treatment over time (Figure 2A-C). The low pH and combined low pH + high T treatments generally followed the trend of increasing oxygen consumption rates over time, with generally higher (low pH) or lower average rates (low pH + high T) compared to the control group (although not significant), whereas the high-T treatment fluctuated around average rates observed in the control group. Respiration rates across the four treatments were significantly impacted by the factor time (*p* = 0.008, Table 3, Figure 3). A significant elevation of oxygen consumption rates averaged over the four treatments (Figure 3) was evident from Week 3 (0.8 µmol O_2_ gDM^-1^ h^-1^) to Week 28 (1.3 µmol O_2_ gDM^-1^ h^-1^) (*p* = 0.014; Table 3). This increase over the course of 25 weeks was followed by a significant decline to 0.68 µmol O_2_ gDM^-1^ h^-1^ over the course of eight weeks from Week 31 to 39 (*p* = 0.007) (Figure 3).

**Figure 2.**
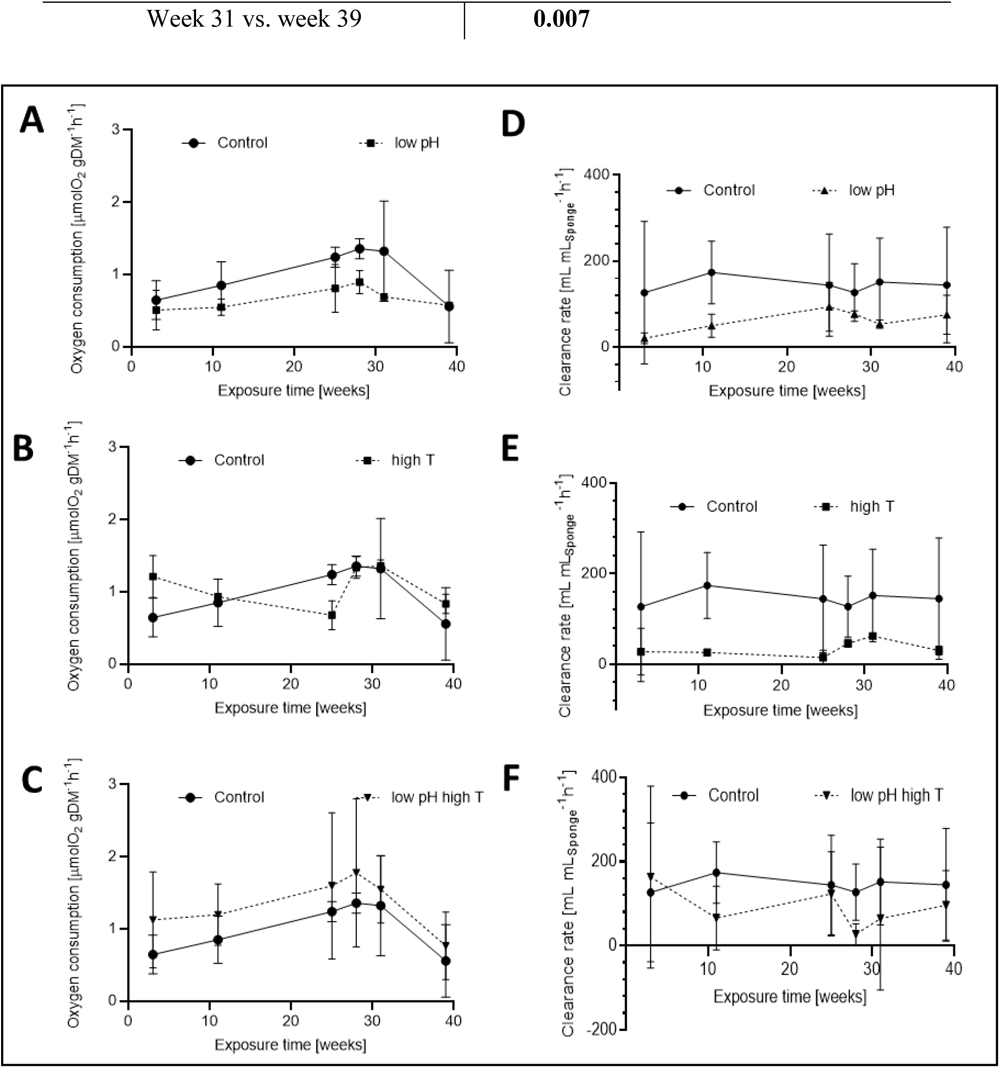
Oxygen consumption (A, B, C) and clearance rates (D, E, F) of *G. barretti* in the control group displayed with rates from the low pH treatment group, high temperature treatment group and low pH + high temperature treatment group over the course of 39 weeks. Values represent the mean of three tanks per treatment and timepoint.

**Figure 3.**
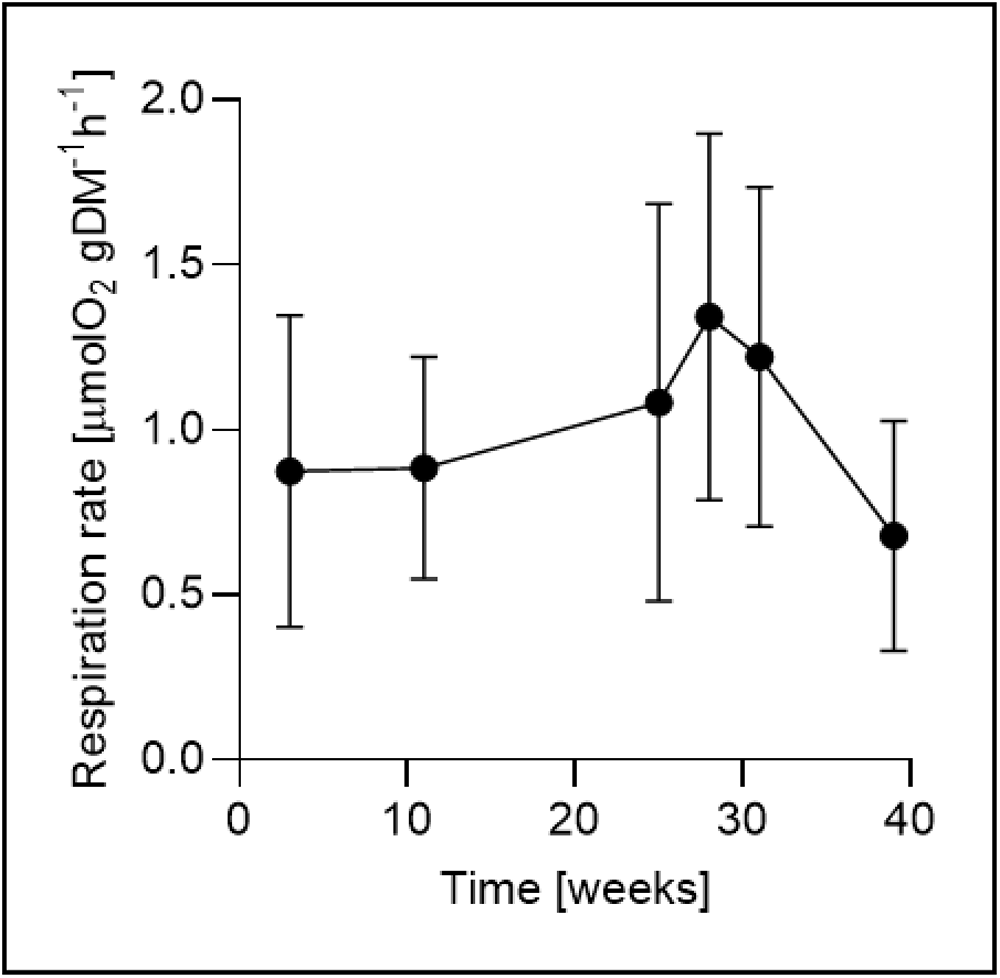
Average of oxygen consumption rates of *G. barretti* across all four treatments over time. Values represent average means of the four treatments per timepoint (n = 12 per time-point). Corresponding letter pairs (ab, cd) indicate a significant difference.

**Table 3.** Overview of the test results for respiration rate over time (Repeated measure mixed effect analysis). For the multiple comparisons, only comparisons with a test result of *p* < 0.05 are stated.

| Fixed factor | *p*-value | F (DFn, DFd) |
| --- | --- | --- |
| Time | **0.008** | F (2.81, 20.84) = 5.293 |
| Temperature | 0.07 | F (1, 8) = 4.11 |
| pH | 0.92 | F (1, 8) = 0.009 |
| Time × Temperature | 0.64 | F (5, 37) = 0.68 |
| Time × pH | 0.76 | F (5, 37) = 0.51 |
| Temperature × pH | 0.12 | F (1,8) = 3.01 |
| Time × Temperature × pH | 0.18 | F (5, 37) = 1.62 |
| **Multiple comparisons (Post-hoc test)** |  |  |
| Week 3 vs. week 28 | **0.014** |  |

### Bacterioplankton clearance rates

Bacterioplankton clearance rates of *G. barretti* were stable throughout time in the control group (*p* = 0.99; Figure 2D). The pH and T treatments followed the same trend over time, with no effect of exposure time on the clearance rate (*p* = 0.98, see Table 4). While the average bacterioplankton clearance rates of the low pH and high T treatment were lower compared to the control treatment, this difference was not significant (*p* = 0.66 for low pH; *p* = 0.21 for high T, Table 4). A significant interaction (*p* = 0.03; Table 4) of high seawater temperature and low pH was detected. This interaction was further explored with a 2-Way ANOVA without the factor “time” but the performed Post-hoc tests did not yield more information about the specifics of the detected interaction (Table 5). Noteworthy, however, is the trend (*p* = 0.09; Table 5) of increased seawater temperature affecting clearance rates in *G. barretti.* This trend is absent in the combined treatments (*p* = 0.64; Table 5) and might point towards an alleviating effect of decreased seawater pH on temperature induced responses.

**Table 4.**
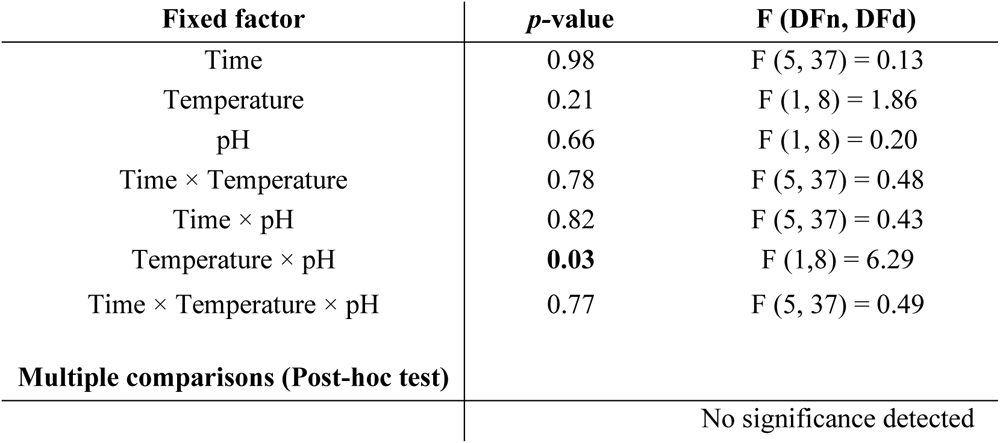
Overview of the test results for clearance rate over time (repeated measure mixed-effect analysis)

**Table 5.** Overview of the two-way ANOVA results for clearance rate with the factors temperature and pH.

| Fixed factor | *p*-value | F (DFn, DFd) |
| --- | --- | --- |
| Temperature | 0.24 | F (1, 8) = 1.86 |
| pH | 0.66 | F (1, 8) = 0.20 |
| Temperature × pH | **0.03** | F (1,8) = 6.29 |
| **Multiple comparisons (Post-hoc test)** |  |  |
| control × high T | 0.09 |  |
| control × low pH | 0.21 |  |
| control × low pH + high T | 0.64 |  |
| high T × low pH + high T | 0.93 |  |
| low pH × low pH + high T | 0.47 |  |
| high T × low pH | 0.78 |  |

### Dissolved inorganic nutrient fluxes

*Geodia barretti* individuals in the control treatment showed a net release of ammonium (NH_4+_), nitrate (NO_3-_), nitrite (NO_2-_) and phosphate (PO_4-_) (Fig. 5; positive value depicts release rates). Fluxes of inorganic nutrients in the pH and T treatments were not significantly affected compared to the control group (*p* > 0.3 in all dissolved inorganic nutrient specific two-way ANOVAs, see details of the tests in the Supplemental Table S1). However, sponges switched from net average release to a net average removal of NO_2-_ in the pH and T treatments, compared to the control.

**Figure 5.**
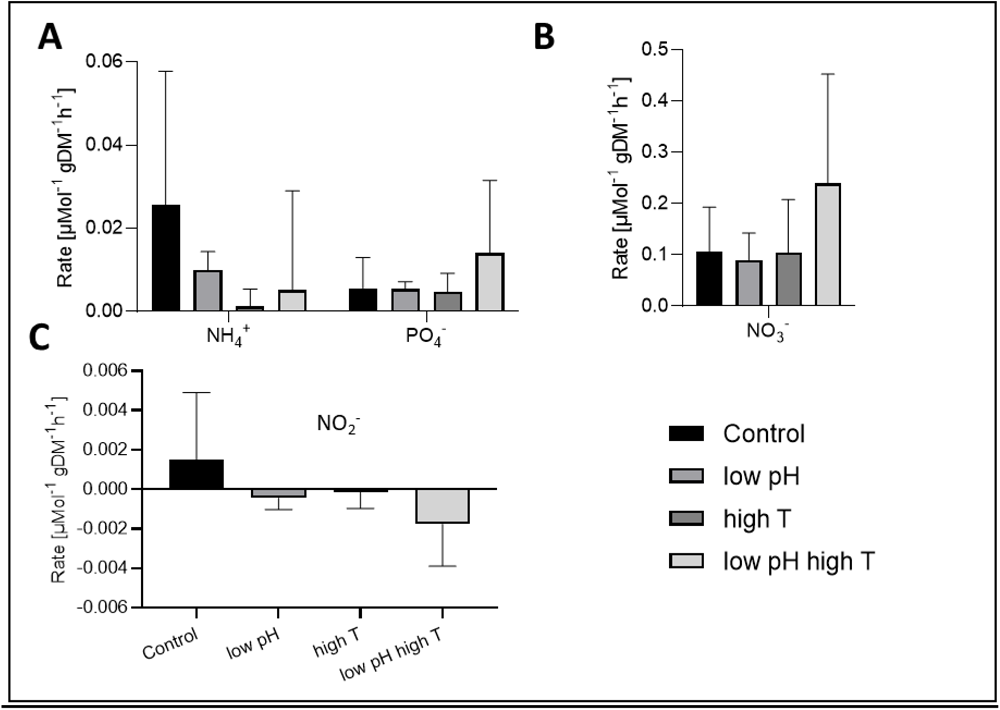
Dissolved inorganic nutrient (A: NH_4+_ and PO_4-_ B: NO_3-_; C: NO_2-_) exchange rates of *G. barretti* in four different treatments at Week 11. Here, positive values represent release, negative values uptake of the respective dissolved inorganic nutrient.

### Tissue necrosis

Throughout the experiment, several individuals from the four treatment groups expressed signs of tissue necrosis. The lowest number of animals showing signs of necrosis was observed in the control group (1 out of 7 over 39 weeks) and the highest number (4 out of 8 over 39 weeks) was observed in the combined treatment group (low pH + high T) (Table 7). However, under the experimental approach, no statistical effect of the treatment on tissue necrosis was evident (*p* = 0.19). Animals found to express signs of necrosis were removed from the experiment.

**Table 7.** Overview of numbers of individuals at the start (n _start_) and the end (n _end_) of the long-term acclimatisation period after 39 weeks. Sponges were removed from the experiment when expressing signs of tissue necrosis.

| **Treatment** | **$n_{\text{start}}$** | **$n_{\text{end}}$** |
| --- | --- | --- |
| control | 7 | 6 |
| low pH | 9 | 6 |
| high T | 7 | 5 |
| low pH high T | 8 | 4 |

## Discussion

Throughout a ten-month *ex situ* study in a flow-through aquarium with *in situ* seawater supply, the metabolic rates (i.e. oxygen consumption, bacterioplankton clearance and dissolved inorganic nutrient removal/release rates) of the deep-sea sponge *G. barretti* were, in general, not significantly affected by simulated future ocean conditions (lower pH, higher temperature). Instead, oxygen consumption rates across all treatments were significantly affected by time of the year with highest rates expressed in late summer. Here, the ecological implications of these findings are discussed, while drawing attention to the complex eco-physiology of the holobiont *G. barretti* and the urgency for holistic approaches to quantify the effects of global changes on this highly abundant deep-sea ecosystem engineer.

### Metabolic rates of G. barretti under present and future pH and T conditions

The measured oxygen consumption rates of individuals in the control treatment of this long-term experiment (0.55-1.36 O_2_ g DM^-1^ h^-1^) were within the range reported for *G. barretti* (1.13‒1.38 O_2_ gDM^-1^ h^-1^) under *ex situ* conditions similar to those in the presented experiment (Strand *et al*., 2017; Leys *et al*., 2018c; Martijn C Bart *et al*., 2020). Biological processes, such as oxygen consumption rates (i.e., as measure for respiration rate) are often driven by the thermodynamics of chemical reactions (Clark and Fraser 2004). This temperature dependency of biological processes is often described by a Q_10_ factor, which represents the magnitude by which the process rate changes with a temperature increase of 10 ⁰C. A Q_10_ value of 2.3 has been reported for respiration in temperate sponges (Riisgard *et al*., 1993; Coma *et al*., 2002). When assuming a similar Q_10_ value for *G. barretti*, the oxygen consumption rates expressed by *G. barretti* exposed to a temperature increase of 4 °C were expected to be elevated by approximately 40 %. However, this was not corroborated by the results from the present experiments, in which significant effects of temperature on oxygen consumption could not be demonstrated. Correspondingly, comparison of average oxygen consumption rates in high temperature treated sponges with rates in control sponges resulted in a Q_10_ value of only 1.18. In contrast, twice higher oxygen consumption rates, corresponding to a Q_10_ value of 6.2, were found in explants of *G. barretti* when exposed to acute heat stress (i.e., a 4‒5 °C elevation in temperature within 24 h) (Strand *et al.,* 2017). Acute exposure to a “heat wave” may trigger acute physiological responses that do not necessarily reflect the long-term acclimatization potential of sponges to elevated temperatures. Despite the more gradual acclimation to higher temperature that was applied in the present study (0.3 °C elevation per 12 h in the present experiment versus 2‒3 °C elevation per 12 h in the experiment by Strand *et al.,* 2017), an initial response was also apparent here, the Q_10_ in Week 3 of the present experiment being 4.8. These findings stress the importance of executing long-term experiments to study acclimation of sponges to future ocean conditions. This is reflected by observations made in two other treatments: The lower pH treatment showed a generally lower average oxygen consumption compared to the control group, but the combined treatment of low pH and high T showed generally higher average oxygen consumption rates. While there is a trend that temperature has an effect (*p* = 0.07) on oxygen consumption rates, the data does not allow for a conclusion if long-term elevated seawater temperatures are causing an increase or decreased of oxygen consumption rates in *G. barretti* under *ex situ* conditions. Changes in oxygen consumption rates can be caused by both favourable (e.g., higher respiration due to elevated nutrition (Ribes et al., 1999)) and unfavourable (e.g., lower respiration due to seizing of pumping (Hoffmann *et al*., 2008) or higher respiration due to higher maintenance costs under suboptimal environmental conditions (Pörtner, Storch and Heilmayer, 2005). Although the manipulated physicochemical parameters were not affecting respiration rates, time of the year was found to have an effect on the oxygen consumption rate in *G. barretti*. Respiration rates were found to significantly increase in late summer. It has been shown that reproduction in fjord populations of *G. barretti* coincides with phytoplankton blooms, gametogenesis/spawning taking place in late summer (Spetland *et al*., 2007). Both reproductive activities and seasonal patterns of environmental conditions have been described to cause fluctuations in physiological rates in sponges (Koopmans and Wijffels, 2008; Koopmans *et al*., 2011; Morley *et al*., 2016). Thus the changes in respiration rates over time in this long-term study could be an indication for reproductive activities in the experimental animals, triggered for example by the influx of biological cues from the wild *G. barretti* fjord population or by changes in abiotic parameters other than temperature and seawater pH occurring outside the location of the experimental facilities. While *ex situ* experiments have the sole purpose to minimize the biological complex environment to controlled conditions, the flow-through design with unfiltered *in situ* deep-sea water was a compromise between keeping sensitive deep-sea sponges alive and the controlled manipulation of physicochemical seawater parameters. One of the environmental variables that was found to vary over time was the abundance of bacterioplankton in the seawater flowing into the experimental facilities. This fluctuation of bacterioplankton indicates that biological and/or chemical *in situ* cues may have affected the sponges in the experimental *ex situ* setup.

Clearance rates for bacterioplankton are another important measure to assess the physiological fitness of filter-feeding sponges (Maldonado, Ribes and van Duyl, 2012). The clearance rates obtained in this experiment (control = 144 ± 58 mL mL_Sponge-1_ h^-1^) are higher than reported for this species (22 ‒ 68 mL mL_Sponge-1_ h^-1^ (Leys *et al.,* 2018). These higher clearance rates could be explained by the size of experimental animals. (de Kluijver *et al*., 2021) have described that smaller individuals of G. barretti assimilate more bacterioplankton than larger individuals. While Leys *et al*. (2018) have used sponges with an average volume of 1 L (maximum 3 L), in this study sponges had an average volume of 0.14 L Similar to the respiration, we found no significant difference of clearance rates between the treatments and the control group. However, clearance rates in the two single stressor treatments were showing a trend of being lower compared to the controls, implying that filter-feeding may be negatively affected by (the combined effects of) pH and T. While this effect was not tested significant, the interaction of pH and high T showed a significant effect. This might imply, that potential effects of single stressors might be alleviated by low pH as clearance rates in the combined treatment largely overlapped with rates observed in the control group. Future studies with higher numbers of replicates are, however, needed to investigate this relation in further details, as the interaction could not be explored to full details with the dataset generated in this experiment.

A third indicator of metabolic rates investigated in this experiment were uptake/release rates of dissolved inorganic nutrients by sponges. *Geodia barretti* is known to exhibit many metabolic nitrogen pathways, mediated by both the host and its microbial symbionts (Leys *et al*., 2018c; Rooks *et al*., 2020; de Kluijver *et al*., 2021). No significant effects could be detected between the control group and the pH and T treatments over time in terms of inorganic nutrient fluxes, indicating no major metabolic changes in the sponges. But changes in average fluxes throughout treatment show interesting trends. In terms of N-cycling, the average release rates of nitrate are increased (i.e. predominantly in the combined pH and T treatment), whereas the fluxes of ammonium and nitrite are decreased, to even a net average uptake of nitrite (Fig. 4). This can imply an increase of aerobic nitrification (i.e., the oxidation of ammonium into nitrite and subsequently nitrate) and/or, but less likely, increased anaerobic processes, such as ammonium oxidation (i.e., loss of nitrogen through transformation of ammonium into nitrite and nitrogen gas). Both processes occur in *G. barretti* and are mediated by the sponge’s microbiome (de Kluijver *et al.,* 2021). Additionally, an increase in the average phosphate (P) release was observed in the combined pH and T treatment compared to the control. Although not conclusive, the increased release of bot N and P indicate an increased assimilation of food under these future scenarios. Unfortunately, we were not able to quantify inorganic nutrient fluxes through time and further studies are needed to relate the fluxes of inorganic nutrients to seasons (e.g., temperature and food availability).

We conclude that the most consistent pattern observed during our 39-week long experiment, namely a summer peak in oxygen consumption, was likely a seasonal response to environmental conditions. Sponge metabolic rates, such as oxygen consumption, but also growth and reproduction, are found to be closely linked with ambient food conditions (Ribes, Coma and Gili, 1999; Spetland *et al*., 2007; Koopmans and Wijffels, 2008; Morley *et al*., 2016). For example, it has been shown that reproduction in fjord populations of *G. barretti* coincides with phytoplankton blooms, gametogenesis/spawning taking place in late summer (Spetland *et al*., 2007). Thus, the recorded physiological changes of the sponges over time in our long-term study may have been caused predominantly by (a)biotic cues in the *in situ* water flowing through the experimental setup.

### Eco-physiology of G. barretti in an Anthropocene ocean

In the North Atlantic Ocean, *G. barretti* has been described to be abundant from depths of 2000 m with temperatures ranging from 3–5 °C until 30 m of water depth in Norwegian fjords ecosystems where individuals can be exposed to seawater temperatures as high as 15 °C (Cárdenas *et al*., 2013). *Geodia barretti’*s distributional range points towards a broad ecological potential. This could explain the continuity of basic physiological parameters across a variety of climate treatments as evidenced in this study and might point towards a resilience of *G. barretti* towards environmental parameters of future NAO conditions. Sponges that contain high abundances of microbial symbionts, such as *G. barretti* (Leys *et al*., 2018c), have been hypothesized to be more resilient to low seawater pH conditions given the diverse, specialized abilities of their microbiome (Goodwin *et al*., 2014). Tissue necrosis is an early warning sign that can result in sponge death, even to *in situ* mass mortality events of *G. barretti* (Guihen *et al.,* 2014). Tissue necrosis has been described as response of sponges to high CO_2_ concentration and increased temperature (Bennett *et al*., 2017; Strand *et al*., 2017) and can be an indication for a disruption of the stable state, or changes in the community composition, of the microbial symbionts associated with the holobiont *G. barretti* (Vacelet and Donadey, 1977; Hentschel *et al*., 2003). At the same time, *G. barretti* heavily relies on the phagocytosis of symbiotic eukaryotes to meet its metabolic carbon requirements (Leys *et al*., 2018c), and the high processing rates of dissolved organic matter (DOM) in *G. barretti* (Bart *et al.,* 2021b) are thought to be partly driven by microbial symbiont activities (Bart et al., 2020a). A shift in the microbial community could therefore affect two energy acquisition pathways in *G. barretti*, limiting its coping potential in a future ocean. While the results of this study show metabolic resilience of *G. barretti* to changing environmental settings, *in situ* observations of mass mortalities following acute temperature increases stress the necessity to acquire *in situ* physiological and metabolic data on deep-sea sponges. Furthermore, first evidence shows that the resilience of *G. barretti* may be overruled when sponges are exposed to multiple stressors, such as re-suspended sediment caused by bottom trawling (Wurz *et al.,* 2021, submitted).

### Limitations of the study

Deep-sea research is a logistically challenging and expensive endeavour (Ruth, 2006), limited by the access to deep-sea organisms and the availability and operation of facilities suitable to maintain them. Despite these limitations, in this experiment, basic physiological parameters in more than 30 deep-sea sponges were assessed. Large fluctuations of physiological rates were evident in the experimental sponges over time. Given this dynamic nature of physiological rates in *G. barretti*, at least five (based on power analysis of here presented data with R wp.anova function) replicates of experimental (tanks) and measurement (sponges) units are advised for future studies, to provide better detection limits for treatment effects. Given the technical challenges to maintain *G. barretti* under *ex situ* conditions, life history traits (age, size, reproductive status, etc.) of the experimental animals most likely differed from natural communities in *G. barretti* dominated sponge grounds (de Kluijver et al. 2021). Here, we used individuals with a diameter of maximum 10 cm and up to 500 g of wet weight. While these smaller-sized sponges are well manageable in the context of aquaria experiments and chamber-based incubations, they deviate from the ambient community structure of *G. barretti* dominated habitats. *Geodia barretti* can reach up to a meter in diameter with a wet weight of up to 24 kg (Klitgaard and Tendal, 2004) and size has recently been found to be a major factor in determining the metabolic rates of sponges (Morganti *et al*., 2019). Furthermore, a recent study suggested that male and female individuals of *Geodia* species spend different amounts of energy on maintenance of reproductive tissues and reproduction (Koutsouveli *et al*., 2020), potentially affecting the gender-specific coping potential towards changing environmental conditions.

### Conclusions

In this *ex situ* study, the deep-sea sponge species *G. barretti* was found to cope with seawater temperature and pH conditions representing future physicochemical parameters in the deep North-Atlantic Ocean. The basic metabolic metrices of oxygen consumption, bacterioplankton clearance and inorganic nutrient exchange rates were not significantly affected by the manipulated seawater temperature and pH treatments. However, the trend towards an effect of elevated seawater temperatures on decreased clearance rates, the observations of potential changes in microbially-mediated nitrogen cycling of *G. barretti*, and the highest level of tissue necrosis in the combined treatment stress the need for further investigation of climate change effects on this deep-sea sponge species. A more holistic approach is needed to further evaluate the eco-physiological resilience of *G. barretti* to future ocean conditions, which should include *in situ* measurements and the broad spectrum of (microbially-mediated) energy acquisition (e.g., the processing of phytoplankton and DOM) processes in *G. barretti*. Care must be taken to include potential interactions between climate change effects and other anthropogenic stressors (e.g., Wurz *et al.,* submitted 2021) when assessing the coping capacity of *G. barretti* towards future ocean conditions.

## Acknowledgements

The Authors thank Heikki and Tone from the University of Bergen for support in maintaining the aquaria facility and technical support when needed. We also thank Detmer Sipkema (WUR) and Rob Joosten (WUR) for generously sharing the flow cytometer and help with analyzing samples.

## References

1. Armstrong, C.W. et al. (2019) ‘Expert assessment of risks posed by climate change and anthropogenic activities to ecosystem services in the deep North Atlantic’, Frontiers in Marine Science [Preprint]. Available at: 10.3389/fmars.2019.00158.

2. Bart, Martijn C. et al. (2020) ‘Differential processing of dissolved and particulate organic matter by deep-sea sponges and their microbial symbionts’, *Scientific Reports* [Preprint]. Available at: 10.1038/s41598-020-74670-0.

3. Bart, Martijn C et al. (2020) ‘Dissolved organic carbon (DOC) is essential to balance the metabolic demands of four dominant North-Atlantic deepsea sponges’, bioRxiv, p. 2020.09.21.305086. Available at: 10.1101/2020.09.21.305086.

4. Bart, M.C. et al. (2021) ‘A Deep-Sea Sponge Loop? Sponges Transfer Dissolved and Particulate Organic Carbon and Nitrogen to Associated Fauna’, Frontiers in Marine Science [Preprint]. Available at: 10.3389/fmars.2021.604879.

5. Beazley, L. et al. (2013) ‘Deep-sea sponge grounds enhance diversity and abundance of epibenthic megafauna in the Northwest Atlantic’, ICES Journal of Marine Science [Preprint]. Available at: 10.1093/icesjms/fst124.

6. Beazley, L. et al. (2018) ‘Predicted distribution of the glass sponge Vazella pourtalesi on the Scotian Shelf and its persistence in the face of climatic variability’, PLoS ONE [Preprint]. Available at: 10.1371/journal.pone.0205505.

7. Bennett, H.M. et al. (2017) ‘Interactive effects of temperature and pCO2 on sponges: from the cradle to the grave’, Global Change Biology [Preprint]. Available at: 10.1111/gcb.13474.

8. Carballo, J. L., & Bell, J. J. (2017). Climate change, ocean acidification and sponges. C.Springer. (2017) Climate Change, Ocean Acidification and Sponges, Climate Change, Ocean Acidification and Sponges. Available at: 10.1007/978-3-319-59008-0.

9. Cárdenas, P. et al. (2013) ‘ Taxonomy, biogeography and DNA barcodes of Geodia species (Porifera, Demospongiae, Tetractinellida) in the Atlantic boreo-arctic region ’, Zoological Journal of the Linnean Society [Preprint]. Available at: 10.1111/zoj.12056.

10. Cárdenas, P. and Rapp, H.T. (2015) ‘Demosponges from the Northern Mid-Atlantic Ridge shed more light on the diversity and biogeography of North Atlantic deep-sea sponges’, Journal of the Marine Biological Association of the United Kingdom [Preprint]. Available at: 10.1017/S0025315415000983.

11. Coma, R. et al. (2002) ‘Seasonality of in situ respiration rate in three temperate benthic suspension feeders’, Limnology and Oceanography [Preprint]. Available at: 10.4319/lo.2002.47.1.0324.

12. Coppari, M., Zanella, C. and Rossi, S. (2019) ‘The importance of coastal gorgonians in the blue carbon budget’, *Scientific Reports* [Preprint]. Available at: 10.1038/s41598-019-49797-4.

13. Danovaro, R., Snelgrove, P.V.R. and Tyler, P. (2014) ‘Challenging the paradigms of deep-sea ecology’, Trends in Ecology and Evolution [Preprint]. Available at: 10.1016/j.tree.2014.06.002.

14. Dijkstra, J.A. et al. (2021) ‘Fine-scale mapping of deep-sea habitat-forming species densities reveals taxonomic specific environmental drivers’, Global Ecology and Biogeography [Preprint]. Available at: 10.1111/geb.13285.

15. De Goeij, J.M. et al. (2013) ‘Surviving in a marine desert: The sponge loop retains resources within coral reefs’, Science [Preprint]. Available at: 10.1126/science.1241981.

15a. De Goeij, J.M., Lesser, M.P. and Pawlik, J.R. (2017) ‘Nutrient fluxes and ecological functions of coral reef sponges in a changing ocean’, in *Climate Change, Ocean Acidification and Sponges: Impacts Across Multiple Levels of Organization*. Available at: 10.1007/978-3-319-59008-0_8.

16. Goodwin, C. et al. (2014) ‘Effects of ocean acidification on sponge communities’, Marine Ecology [Preprint]. Available at: 10.1111/maec.12093.

17. Guihen, D., White, M. and Lundälv, T. (2014) ‘Temperature shocks and ecological implications at a cold-water coral reef’, ANZIAM Journal [Preprint]. Available at: 10.1017/S1755267212000413.

18. Hanz, U. et al. (2021) ‘Long-term Observations Reveal Environmental Conditions and Food Supply Mechanisms at an Arctic Deep-Sea Sponge Ground’, Journal of Geophysical Research: Oceans [Preprint]. Available at: 10.1029/2020jc016776.

19. Hawkes, N. et al. (2019) ‘Glass sponge grounds on the Scotian Shelf and their associated biodiversity’, Marine Ecology Progress Series [Preprint]. Available at: 10.3354/meps12903.

20. Hentschel, U. et al. (2003) ‘Microbial diversity of marine sponges.’, Progress in molecular and subcellular biology [Preprint]. Available at: 10.1007/978-3-642-55519-0_3.

21. Hoffmann, F. et al. (2008) ‘Oxygen dynamics and transport in the Mediterranean sponge Aplysina aerophoba’, Marine Biology [Preprint]. Available at: 10.1007/s00227-008-0905-3.

22. Hogg, M.M. et al. (2010) ‘Deep Sea Sponge Grounds: Reservoirs of Biodiversity’, in *UNEP-WCMC Biodiversity Series*. Available at: 10.1016/S0378-777X(80)80057-6.

23. Kazanidis, G. et al. (2018) ‘Unravelling the versatile feeding and metabolic strategies of the cold-water ecosystem engineer Spongosorites coralliophaga (Stephens, 1915)’, Deep-Sea Research Part I: Oceanographic Research Papers [Preprint]. Available at: 10.1016/j.dsr.2018.07.009.

24. Kazanidis, G. et al. (2019) ‘Distribution of deep-sea sponge aggregations in an area of multisectoral activities and changing oceanic conditions’, Frontiers in Marine Science [Preprint]. Available at: 10.3389/fmars.2019.00163.

25. Kenchington, E., Power, D. and Koen-Alonso, M. (2013) ‘Associations of demersal fish with sponge grounds on the continental slopes of the northwest Atlantic’, Marine Ecology Progress Series [Preprint]. Available at: 10.3354/meps10127.

26. Klitgaard, A.B. and Tendal, O.S. (2004) ‘Distribution and species composition of mass occurrences of large-sized sponges in the northeast Atlantic’, Progress in Oceanography [Preprint]. Available at: 10.1016/j.pocean.2004.06.002.

27. de Kluijver, A. et al. (2021) ‘An Integrative Model of Carbon and Nitrogen Metabolism in a Common Deep-Sea Sponge (Geodia barretti)’, Frontiers in Marine Science [Preprint]. Available at: 10.3389/fmars.2020.596251.

28. Koopmans, M. et al. (2011) ‘Carbon conversion and metabolic rate in two marine sponges’, Marine Biology [Preprint]. Available at: 10.1007/s00227-010-1538-x.

29. Koopmans, M. and Wijffels, R.H. (2008) ‘Seasonal growth rate of the sponge Haliclona oculata (Demospongiae: Haplosclerida)’, *Marine Biotechnology* [Preprint]. Available at: 10.1007/s10126-008-9086-9.

30. Koutsouveli, V. et al. (2020) ‘Reproductive Biology of Geodia Species (Porifera, Tetractinellida) From Boreo-Arctic North-Atlantic Deep-Sea Sponge Grounds’, Frontiers in Marine Science [Preprint]. Available at: 10.3389/fmars.2020.595267.

31. Kutti, T., Bannister, R.J. and Fosså, J.H. (2013) ‘Community structure and ecological function of deep-water sponge grounds in the Traenadypet MPA-Northern Norwegian continental shelf’, Continental Shelf Research [Preprint]. Available at: 10.1016/j.csr.2013.09.011.

32. Leys, S.P. et al. (2018a) ‘Phagocytosis of microbial symbionts balances the carbon and nitrogen budget for the deep-water boreal sponge Geodia barretti’, Limnology and Oceanography [Preprint]. Available at: 10.1002/lno.10623.

33. Leys, S.P. et al. (2018b) ‘Phagocytosis of microbial symbionts balances the carbon and nitrogen budget for the deep-water boreal sponge Geodia barretti’, Limnology and Oceanography [Preprint]. Available at: 10.1002/lno.10623.

34. Leys, S.P. et al. (2018c) ‘Phagocytosis of microbial symbionts balances the carbon and nitrogen budget for the deep-water boreal sponge Geodia barretti’, Limnology and Oceanography [Preprint]. Available at: 10.1002/lno.10623.

35. Luter, H.M. et al. (2017) ‘Microbiome analysis of a disease affecting the deep-sea sponge Geodia barretti’, FEMS microbiology ecology [Preprint]. Available at: 10.1093/femsec/fix074.

36. Maldonado, M., Ribes, M. and van Duyl, F.C. (2012) ‘Nutrient Fluxes Through Sponges. Biology, Budgets, and Ecological Implications’, in Advances in Marine Biology. Available at: 10.1016/B978-0-12-394283-8.00003-5.

37. Morganti, T.M. et al. (2019) ‘Size Is the Major Determinant of Pumping Rates in Marine Sponges’, Frontiers in Physiology [Preprint]. Available at: 10.3389/fphys.2019.01474.

38. Morley, S.A. et al. (2016) ‘Extreme phenotypic plasticity in metabolic physiology of Antarctic demosponges’, Frontiers in Ecology and Evolution [Preprint]. Available at: 10.3389/fevo.2015.00157.

39. Ottaviani, D. (2020) Economic valuation of ecosystem services provided by deep-sea sponges. Vol. 1217., Economic valuation of ecosystem services provided by deep-sea sponges. Vol. 1217. Available at: 10.4060/cb2331en.

40. Pörtner, H.O., Storch, D. and Heilmayer, O. (2005) ‘Constraints and trade-offs in climate-dependent adaptation: Energy budgets and growth in a latitudinal cline’, Scientia Marina [Preprint]. Available at: 10.3989/scimar.2005.69s2271.

41. Ribes, M., Coma, R. and Gili, J.M. (1999) ‘Natural diet and grazing rate of the temperate sponge Dysidea avara (Demospongiae, Dendroceratida) throughout an annual cycle’, Marine Ecology Progress Series [Preprint]. Available at: 10.3354/meps176179.

42. Riebesell, U., Fabry, V.J. and Hansson, L. (2010) ‘Guide to best practices for ocean acidification research and data reporting’, European commission [Preprint].

43. Riisgard, H.U. et al. (1993) ‘Suspension feeding in marine sponges Halichondria panicea and Haliclona urceolus: effects of temperature on filtration rate and energy cost of pumping’, Marine Ecology Progress Series [Preprint]. Available at: 10.3354/meps096177.

45. Rix, L. et al. (2016) ‘Coral mucus fuels the sponge loop in warm-and cold-water coral reef ecosystems’, *Scientific Reports* [Preprint]. Available at: 10.1038/srep18715.

46. Roberts, E. et al. (2021) ‘Water masses constrain the distribution of deep-sea sponges in the North Atlantic Ocean and Nordic Seas’, Marine Ecology Progress Series [Preprint]. Available at: 10.3354/meps13570.

47. Robertson, L.M., Hamel, J.F. and Mercier, A. (2017) ‘Feeding in deep-sea demosponges: Influence of abiotic and biotic factors’, Deep-Sea Research Part I: Oceanographic Research Papers [Preprint]. Available at: 10.1016/j.dsr.2017.07.006.

48. Rooks, C. et al. (2020) ‘Deep-sea sponge grounds as nutrient sinks: Denitrification is common in boreo-Arctic sponges’, Biogeosciences [Preprint]. Available at: 10.5194/bg-17-1231-2020.

49. Rossi, S. and Rizzo, L. (2020) ‘Marine Animal Forests as Carbon Immobilizers or Why We Should Preserve These Three-Dimensional Alive Structures’, in *Perspectives on the Marine Animal Forests of the World*. Available at: 10.1007/978-3-030-57054-5_11.

50. Ruth, L. (2006) ‘Gambling in the deep sea’, *EMBO Reports* [Preprint]. Available at: 10.1038/sj.embor.7400609.

50a. Spetland, F. et al. (2007) ‘Sexual reproduction of Geodia barretti Bowerbank, 1858 (Porifera, Astrophorida) in two Scandinavian fjords’, Museu Nacional Serie Livros [Preprint].

51. Stevenson, A. et al. (2020) ‘Warming and acidification threaten glass sponge Aphrocallistes vastus pumping and reef formation’, *Scientific Reports* [Preprint]. Available at: 10.1038/s41598-020-65220-9.

52. Strand, R. et al. (2017) ‘The response of a boreal deep-sea sponge holobiont to acute thermal stress’, *Scientific Reports* [Preprint]. Available at: 10.1038/s41598-017-01091-x.

53. Sweetman, A.K. et al. (2017) ‘Major impacts of climate change on deep-sea benthic ecosystems’, Elementa [Preprint]. Available at: 10.1525/elementa.203.

54. Tjensvoll, I. et al. (2013) ‘Rapid respiratory responses of the deep-water sponge Geodia barretti exposed to suspended sediments’, Aquatic Biology [Preprint]. Available at: 10.3354/ab00522.

55. Vacelet, J. and Donadey, C. (1977) ‘Electron microscope study of the association between some sponges and bacteria’, Journal of Experimental Marine Biology and Ecology [Preprint]. Available at: 10.1016/0022-0981(77)90038-7.

